# Rod signals are routed through specific Off cone bipolar cells in primate retina

**DOI:** 10.1101/2020.05.17.100982

**Authors:** Amanda J. McLaughlin, Kumiko A. Percival, Jacqueline Gayet-Primo, Teresa Puthussery

## Abstract

Adapting between scotopic and photopic illumination involves switching the routing of retinal signals between rod and cone-dominated circuits. In the daytime, cone signals pass through parallel On and Off cone bipolar cells, that are sensitive to increments and decrements in luminance, respectively. At night, rod signals are routed into these cone-pathways via a key glycinergic interneuron, the AII amacrine cell (AII-AC). In primates, it is not known whether AII-ACs contact all Off-bipolar cell types indiscriminately, or whether their outputs are biased towards specific Off-bipolar cell types. Here, we show that the rod-driven glycinergic output of AII-ACs is strongly biased towards a subset of macaque Off-cone bipolar cells. The Off-bipolar types that receive this glycinergic input have sustained physiological properties and include the Off-midget bipolar cells, which provide excitatory input to the Off-midget ganglion cells (parvocellular pathway). The kinetics of the glycinergic events are consistent with the involvement of the α1 glycine receptor subunit. Taken together with results in mouse retina, our findings point towards a conserved motif whereby rod signals are preferentially routed into sustained Off signaling pathways.

**Significance Statement:** Visual signals pass through different retinal neurons depending on the prevailing level of illumination. Under night-time light levels, signals from rods pass through the AII amacrine cell, an inhibitory interneuron that routes rod signals into On and Off bipolar cells to detect increments and decrements in light intensity, respectively. Here, we show in primate retina that the output of AII amacrine cells is strongly biased towards specific Off bipolar cell types, which suggests that rod signals reach the brain via specific neural channels. Our results further our understanding of how visual signals are routed through visual circuits during night-time vision.

## Introduction

Understanding human retinal function requires knowledge of the synaptic and circuit mechanisms that are engaged under different illumination levels. Signals from rod photoreceptors are routed through different circuits depending on illumination level (Müller et al., 1988; Grimes et al., 2018b). Under scotopic (night-time) light levels, the primary rod pathway transmits signals from rods to rod bipolar cells, which in turn signal to AII amacrine cells (AII-ACs). The AII-ACs then relay signals to Off-cone bipolar cells (CBCs) through sign-inverting glycinergic synapses, and to On CBCs through sign-conserving gap-junctions (Kolb and Famiglietti, 1974; Famiglietti and Kolb, 1975; Bloomfield and Dacheux, 2001). In this way, increments and decrements in light intensity detected by the rods, and collected by the On-type rod-bipolar cells, are routed appropriately into the On and Off pathways. As light levels rise into the mesopic (twilight) range, rod signals pass to cones via gap junctions (secondary rod pathway) and are then transmitted through the CBC pathways (Nelson, 1977; Schneeweis and Schnapf, 1995, 1999). A tertiary rod pathway also operates at mesopic light levels, in which rods synapse directly with Off-CBCs (Soucy et al., 1998; Hack et al., 1999; Tsukamoto et al., 2001; Li et al., 2010).

A recent study showed that primates differ from other species in that rod signals are routed principally through the primary rod pathway, via AII-ACs, even at mesopic light levels (Grimes et al., 2018a). However, it is not known whether all of the five primate Off-CBC types (Boycott and Wässle, 1991; Haverkamp et al., 2003; Puthussery et al., 2013, 2014; Peng et al., 2019) receive AII -AC input, and thus participate in primary rod pathway signaling. Ultrastructural reconstructions in mice show that the majority of glycinergic output synapses from AII-ACs are directed towards a single Off-CBC subtype (CBC2) (Tsukamoto and Omi, 2017; Graydon et al., 2018; Gamlin et al., 2020). If glycinergic inhibition from primate AII-ACs is similarly biased towards specific Off-CBC types, the temporal properties of rod signals would be shaped by these bipolar cells.

Routing of signals from AII-AC to Off-CBCs also occurs under photopic light levels, when rods are saturated. Under these conditions, cone activation tonically depolarizes the On-CBCs, which in turn depolarize the AII-ACs via gap junctions such that they increase their glycinergic output onto Off-CBC axon terminals (Liang and Freed, 2010). Such “crossover” inhibition from the On pathway hyperpolarizes the Off-CBCs below threshold for glutamate release and thus rectifies their output (Liang and Freed, 2010). Understanding AII-AC to Off-CBC connectivity is thus also important for understanding signal routing under photopic conditions.

Here, we use a functional approach to determine whether the glycinergic output from macaque AII-ACs is directed towards specific Off-CBC types. We find evidence that primate AII-AC signals are strongly biased towards Off-CBCs that have sustained physiological response properties. Our results suggest there may be heterogeneity in the scotopic sensitivity and linearity of primate Off-CBCs and, by extension, their postsynaptic ganglion cells.

## Materials & Methods

### Tissue preparation

Eyes from adult rhesus macaques (*Macaca mulatta)* of either sex were collected from animals euthanized for unrelated projects at the Oregon or California National Primate Research Centers. Eyes were enucleated immediately *post-mortem*, the anterior eye and vitreous were removed, and posterior eyecups were transported in bicarbonate-buffered Ames’ medium (U.S. Biologicals) equilibrated with 95% O_2_/5% CO_2_ at 22-25°C (pH=7.4). After ∼1hr, the retina/RPE/choroid complex was dissected free of the sclera and maintained in the same buffer until slices were made. For slice preparation, the neural retina was isolated from the RPE/choroid and ∼300 μm thick slices were made using a vibrating blade microtome or tissue chopper. For vibratome sections, retinas were embedded in 3% low-melting point agar in HEPES-Ames’ (pH=7.4). All samples were from peripheral (>8 mm) superonasal or inferonasal retina. Slices were prepared and recordings made under low photopic background illumination levels.

### Immunohistochemistry

Macaque retinae were briefly perfusion- or immersion-fixed in 4% paraformaldehyde in 0.1M PB or PBS for 30-60 mins. After fixation, tissues were rinsed in PBS, cryoprotected in graded sucrose solutions (10%, 20%, 30%) and stored at -20°C until use. Retinal pieces were embedded in Cryo-gel embedding medium and cryosectioned at 14-16 μm. For immunohistochemistry, sections were blocked in 10% normal horse serum, 0.3-1% Triton X-100, 0.025% NaN_3_ in PBS for an hour. Primary antibodies were diluted in 3% normal horse serum, 0.3-1% Triton X-100, 0.025% NaN_3_ in PBS and applied overnight at 20-25°C. The primary antibodies were: ms anti-Kir2.1 (Clone N112B/14, Neuromab, Cat. # 73-210, tissue culture supernatant, RRID: AB_11001668, dilution 1:15), ms anti-Ca_V_3.1 (Clone N178A/9; Cat. # Neuromab, 73-206, tissue culture supernatant, RRID: AB_10673097, dilution 1:10), sheep anti-secretagogin (BioVendor, RD1884120100, RRID: AB_2034060, dilution 1:2000), rabbit anti-GLT-1 (Tocris, #2063, dilution 1:2000). Secondary antibodies against mouse, rabbit or sheep antigens were raised in donkey and conjugated to Alexa 488 or Alexa 594 (Molecular Probes, ThermoFisher, RRIDs: AB_141607, AB_2534083, AB_141637). Samples were mounted in Mowiol. Confocal images were acquired on an Olympus FV1000 (60/1.4 objective, 488, 559, and 643 nm laser lines) or a Zeiss LSM 880 laser scanning microscope (63/1.4 objective, 488, 561, and 594 nm laser lines). Images were acquired sequentially to prevent fluorescent cross talk. In some cases, adjustments to image brightness and contrast were made with ImageJ (Fiji).

### Bipolar cell electrophysiology

For whole-cell patch-clamp recordings, retinas were superfused with warm (31-33°C) bicarbonate-buffered Ames’ medium at ∼2-3 ml/min. Patch electrodes (9-12 MΩ) were coated in Parafilm to reduce pipette capacitance. Electrodes were filled with an intracellular solution containing (in mM): 130 K-methanesulfonic acid, 8 KCl, 2 Mg_2_-ATP, 1 Na-GTP, 1 EGTA, 10 Na_0.5_-HEPES and ∼0.1 Alexa Fluor 488 hydrazide adjusted to pH 7.35 with KOH (osmolarity = 290 mOsm). In some cases, 10 mM phospho-creatine was included in the intracellular solution. To assess recovery from inactivation for T-type calcium currents, a Cs^+^ based intracellular solution was used to block voltage-gated potassium current. This solution contained (in mM): 120 Cs-methanesulfonic acid, 8 KCl, 2 Mg_2_-ATP, 1 Na-GTP, 1 EGTA, 10 Na_0.5_-HEPES, 10 mM phosophocreatine and ∼0.1 Alexa Fluor 488 hydrazide adjusted to pH 7.35 with CsOH (osmolarity = 290 mOsm). Reported voltages were corrected for a liquid junction potential of -10 mV unless otherwise indicated. Currents were filtered at a -3 dB cut-off frequency of 2 kHz by the 4 pole Bessel filter of a HEKA EPC-10 double patch amplifier, and digitized at 10 kHz. Series resistance was compensated online and monitored during recordings. Cells were excluded from analysis if series resistance was greater than 35 MΩ.

For pharmacological experiments, stock solutions were made in ultrapure water and aliquots stored at -20°C until further use. GYKI-53655, L-AP4 and tetrodotoxin were obtained from Tocris Bioscience and strychnine was obtained from Sigma. For all pharmacological experiments, drugs were superfused for at least 3 mins to ensure that steady-state drug effects had been reached. CPPG (Tocris Bioscience) was diluted in HEPES-buffered Ames’ medium and focally applied to the outer plexiform layer via a ∼5 MΩ patch pipette. For these experiments, bath flow was directed towards the outer retina to obviate drug effects on the inner retinal circuitry.

### Data analysis

#### Current variance

The current variance measurements in Fig. 1 were obtained during a voltage-step to - 45 mV. To exclude contributions from time-varying voltage-gated current components (e.g. A-type potassium currents), measurements were made after voltage-gated currents had decayed to a steady-state. For variance measurements before and after puff application of CPPG (Fig. 4), a 0.5 s time window was analyzed before and after drug application.

**Figure 1.**
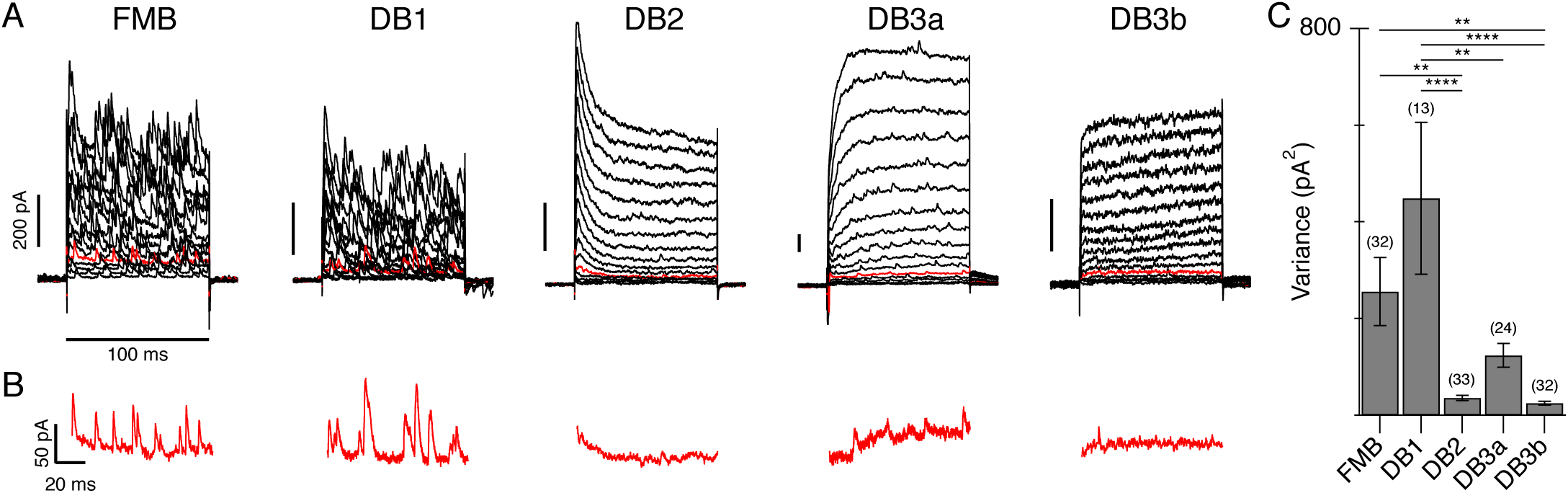
Tonic inhibition is present in specific Off-CBC types. ***A***, Voltage-activated currents in representative FMB, DB1, DB2, DB3a and DB3b cells. The stimulus was a series of depolarizing voltage steps from a holding potential of -70 mV (+5 mV increments, 16 steps). The red traces are the currents at a step potential of -45 mV, which are also shown on an expanded timescale in ***B***. Note the greater number of spontaneous events in FMB and DB1 cells compared to other cell types. Traces in ***B*** show the currents from 5 ms after the onset of the voltage step to the end of the voltage step. ***C***, Bar graph showing current variance in the different Off-CBCs during a voltage step to - 45 mV. Number of cells for each type is indicated above bars. Data are mean±s.e.m. ** p < 0.01, **** p < 0.0001 with Tukey’s post-test.

**Figure 2.**
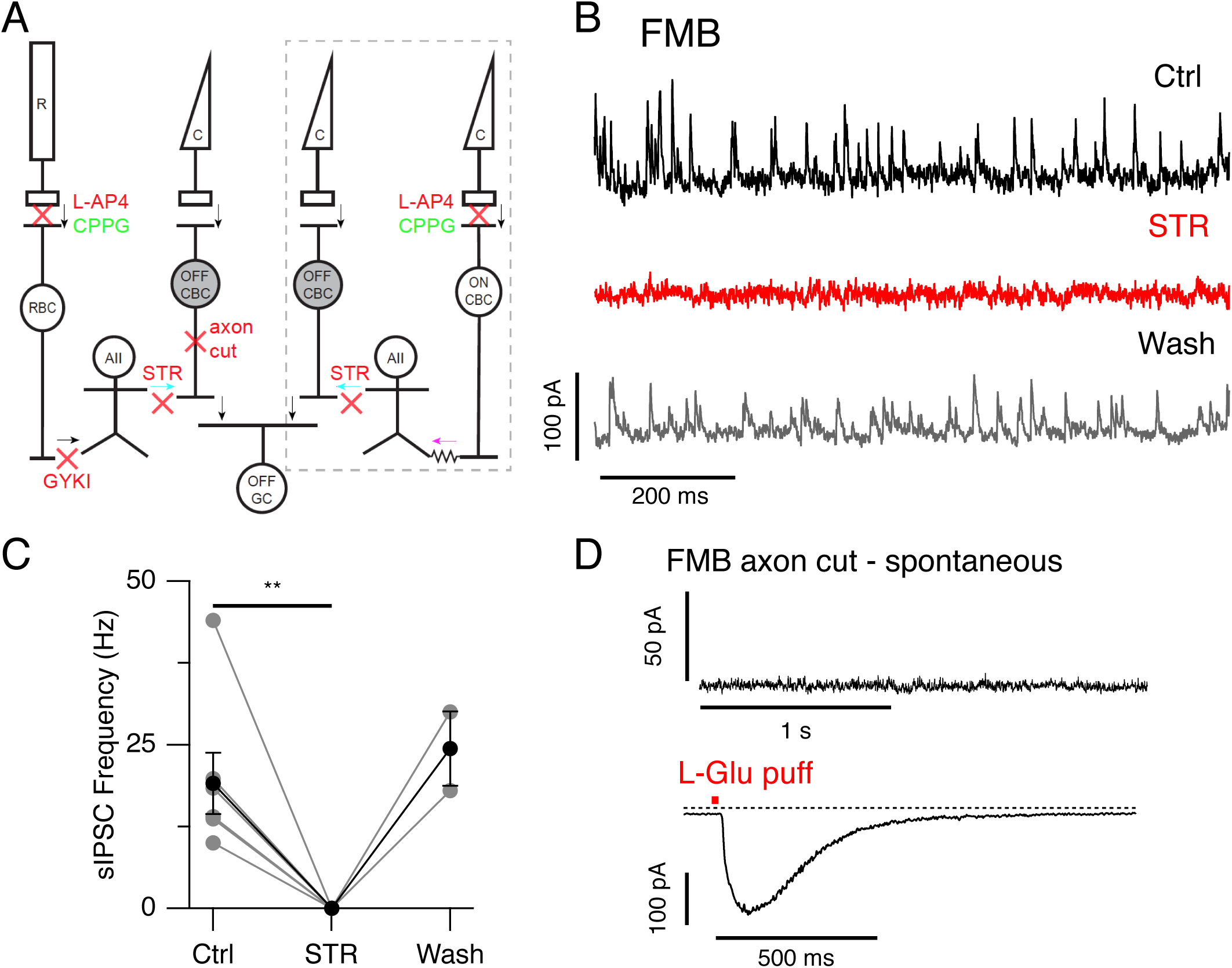
Tonic inhibition in FMB cells is glycinergic and arises at the axon terminals. ***A***, Schematic showing glycinergic inhibition onto Off-CBCs through rod and cone pathways. Arrows indicate direction of signal flow and red crosses indicate site of action of pharmacological agents used herein. Under scotopic conditions, light signals from rods are transmitted to rod bipolar cells (RBC) via mGluR6 receptors. This synapse can be blocked with L-AP4 and activated with CPPG. RBCs transmit signals to AII-ACs (AII) via GYKI-sensitive AMPA receptors. AII-ACs inhibit Off-CBCs through glycinergic synapses (blue arrows) that can be blocked with strychnine (STR). Under photopic conditions (grey dotted box), cones relay light increments to On-CBCs via mGluR6 receptors (On CBC). On-CBCs drive AII-ACs via gap junctions (magenta arrow) and AII-ACs in turn inhibit Off-CBCs via glycinergic synapses. In both pathways, Off-CBCs are relayed to Off-ganglion cells (OFF GCs). ***B***, Example traces from an FMB cell showing the frequency of sIPSCs before, during and after bath application of strychnine (0.5 μM). Events were recorded during a voltage step to -20 mV from a holding potential of -70 mV. ***C***, Effect of STR on sIPSC frequency for a group of FMB cells (Ctrl & STR; n=7 cells, washout; n=2 cells, **p=0.004, Wilcoxon signed rank test). Gray points indicate individual cells, black points indicate mean±s.e.m. ***D***, Top panel, example trace showing a lack of spontaneous events in an axotomized FMB cell during a voltage step to -20 mV from -70 mV. Bottom panel, current response to a 20 ms puff application (red bar) of L-glutamate (1 mM) applied to the same cell. The presence of a glutamate-evoked inward current confirms the Off-type physiology of the axotomized midget cell. Holding potential is -70 mV.

**Figure 3.**
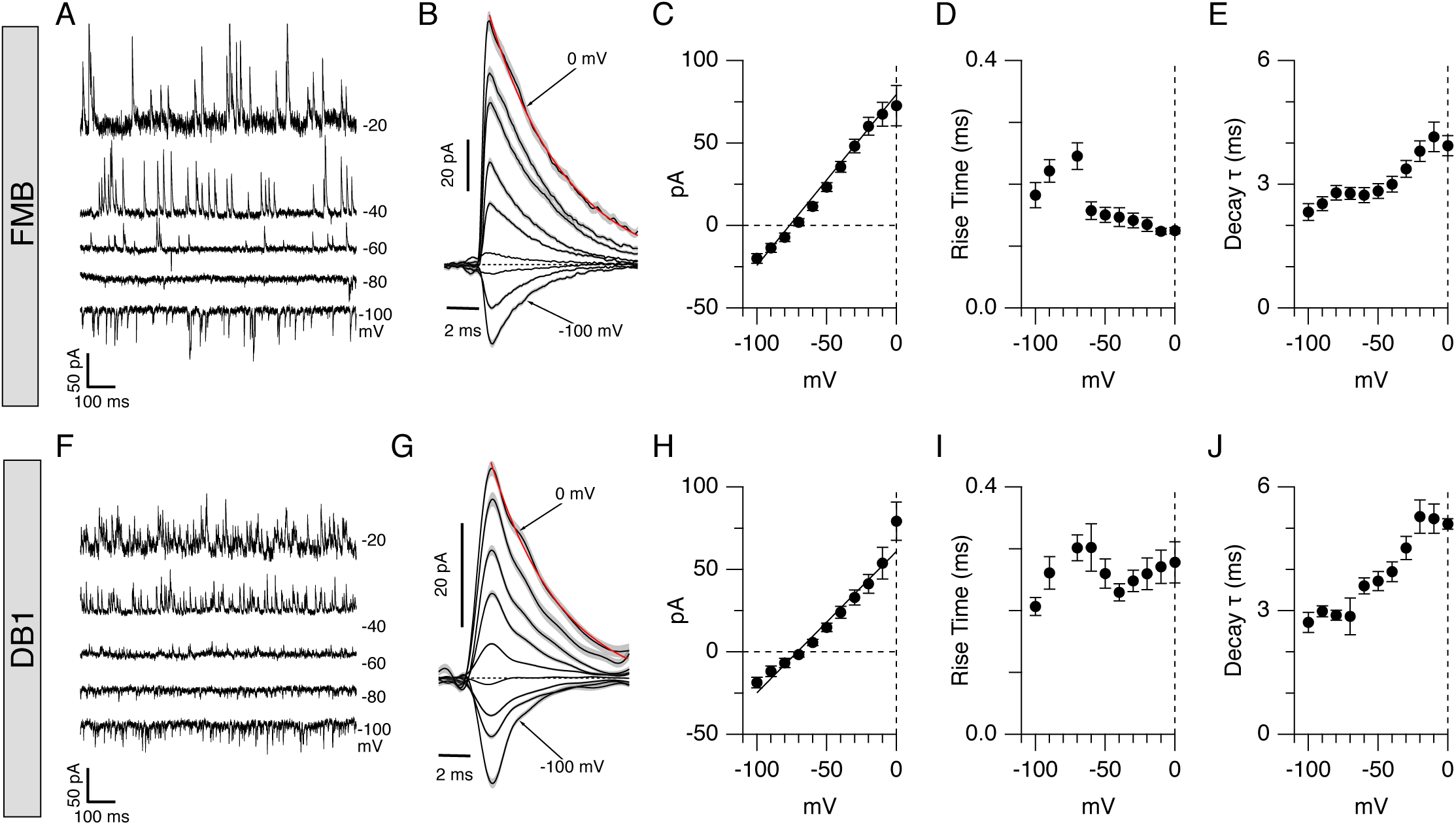
Properties of sIPSCs in FMB and DB1 bipolar cells. Example traces and quantification of sIPSC events in FMB (**A-E**) and DB1 (**F-J**) bipolar cells. ***A***, Example traces showing sIPSCs in an FMB cell at the holding potentials indicated to the right of traces. ***B***. Average sIPSC events extracted from the same bipolar cell shown in ***A*** using a matched template filter (average of 64-126 events per voltage). Gray shading shows s.e.m. Red trace shows single exponential fit to event decay at 0 mV. ***C***, Current-voltage plot showing sIPSC amplitude as a function of voltage for a group of FMB cells (n =20 cells). Note that events reverse close to the inhibitory reversal potential. ***D-E***, Average sIPSC rise-time (***D***) and decay time constants (***E***) for a group of FMB cells. ***F***, Example traces showing sIPSCs in a DB1 cell at holding potentials indicated to right of traces. ***G***. Average sIPSC events from a single DB1 cell held at different voltages (average of 70-130 events per voltage). These events are from the same cell shown in A. Gray shading shows s.e.m. Red trace shows single exponential fit to event decay at 0 mV. ***H***, Current-voltage plot of average sIPSC events for a group of DB1 cells (n=7 cells). ***I-J***, Average sIPSC rise-time (***I***) and decay time constants (***J***) from a group of FMB cells. All graphs show mean ± s.e.m.

**Figure 4.**
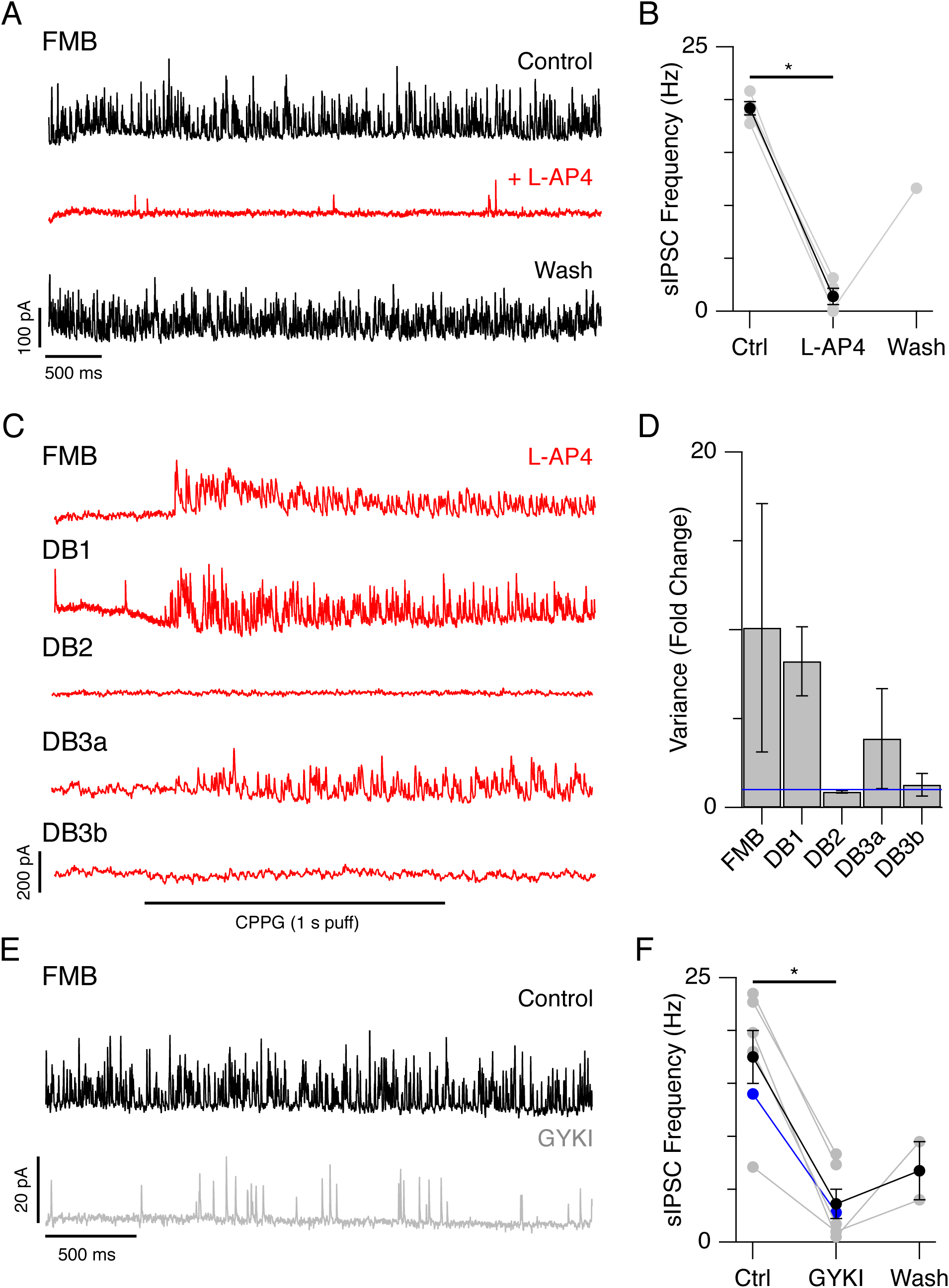
sIPSCs are driven by RBC input to AII amacrine cells. ***A-B***, Example traces showing sIPSCs in an FMB cell before, during and after bath-application of L-AP4 (10 μM). ***B***. Effect of L-AP4 on sIPSC frequency for a group of FMB cells (Ctrl & L-AP4; n=4 cells, washout; n=1 cell, *p = 0.028, Wilcoxon test). Gray points represent individual cells, black points show mean±s.e.m. ***C***, Example traces showing effect of puffing CPPG (600 μM) in the OPL in the presence of bath-applied L-AP4. Note that CPPG puffs increase the frequency of sIPSCs in FMB, DB1 and DB3a cells but not in DB2 and DB3b cells. ***D***, Bar graph showing the fold change in current variance before and after puff-application of CPPG in the presence of L-AP4. Blue line represents y=1. Data are mean ± s.e.m. from n=3 FMB, n=2 DB1, n=3 DB2, n=2 DB3a and n=2 DB3b cells. Current traces in A and C are from a step potential of -20 mV. ***E-F***, Example traces showing reduced frequency of sIPSCs in an FMB cell after bath-application of the AMPA receptor antagonist, GYKI 53655 (10 μM). Holding potential is - 60 mV. ***F***. Effect of GYKI on sIPSC frequency for a group of 5 FMB cells (gray points) and 1 DB1 cell (blue point). Partial washout was obtained in n=2 FMB cells. *p = 0.016 with Wilcoxon rank test. Black points show mean±s.e.m.

#### IPSC analysis and pharmacology

sIPSCs were detected and extracted from recordings using a matched template filter approach implemented in Igor Pro 8. First, average event templates were obtained using an amplitude threshold and average event rise and decay times. A matched template filter was then used to convolve the data traces with the templates. 10-90% rise times were measured from a sigmoidal fit and IPSC decay time constants were determined using a single exponential fit. For quantification of the effects of strychnine, L-AP4 and GYKI 53655 on sIPSCs, IPSC event frequency was estimated using an amplitude threshold cut-off of two standard deviations above the mean current level. Only events with a minimum width of 1 ms were counted to prevent detection of non-neural noise.

#### Voltage-gated currents

To obtain the net voltage-activated component of the membrane current, scaled linear current components were subtracted from the raw recorded currents. To obtain empirical estimates of the activation range and voltage-dependence of voltage-gated currents, leak-subtracted current-voltage (I-V) relations were fit using a Boltzmann equation, and assuming ohmic conductance:

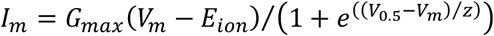

where, *G*_*max*_ is the maximum conductance, *E*_*ion*_ is the reversal potential, *V*_*0.5*_ is the half-activation potential, and *z* is the voltage-sensitivity. All analysis was performed with custom routines in Igor Pro 7 or 8 (Wavemetrics, Lake Oswego, OR).

The time constant of T-type current activation was measured by fitting a sigmoid to the activation phase and calculating the 10-90% rise time. The inactivation time-course was assessed by fitting the decay of the current waveforms at the peak potential of the I-V relationships with a single exponential function. Recovery from inactivation for T-type currents was measured using a paired-pulse protocol. Cells were depolarized with a 500 ms pre-pulse from -90 mV to -40 mV and a second test pulse (−90 to - 40 mV, 100 ms) was given at increasing time intervals after the pre-pulse. Recovery was expressed as the amplitude of the test pulse as a fraction of the pre-pulse amplitude. The time constant of recovery from inactivation was measured by fitting a double-exponential function where *τ*_1_ and *τ*_2_ represent a fast and slow component of recovery.

#### Visualizations of single-cell RNA sequencing data

Mining and visualizations of single-cell RNA-sequencing expression profiles from peripheral primate bipolar cells (Peng et al., 2019) were generated using the Single Cell Portal, (https://singlecell.broadinstitute.org/single_cell).

#### Experimental Design & statistical analysis

Statistical comparisons were made using one-way ANOVA, unpaired t-tests or the Wilcoxon signed-rank test as indicated. For each experiment, the number of cells is indicated in the text or in the figure legends. Data was tested for normality and Welches correction was applied when variances were unequal. An alpha level of 0.05 was used for statistical significance. All statistical analyses were performed in Igor Pro 8.0.

## Results

### Prominent spontaneous inhibitory input in a subset of Off-CBC types

Our objective was to determine whether glycinergic output synapses from AII-ACs are preferentially directed towards specific Off-CBC types. There are five Off-CBC types in macaque and human retina, which differ with respect to their morphology, transcriptomic profiles and functional properties (Boycott and Wässle, 1991; Haverkamp et al., 2003; Puthussery et al., 2013, 2014; Peng et al., 2019). We first compared the frequency of spontaneous inhibitory postsynaptic currents (IPSCs) in the different Off-CBC types that were distinguished based on their morphology and inventory of voltage-gated channels ((Puthussery et al., 2013) and see Fig 7). Figure 1 shows representative examples of voltage-activated currents in each Off-CBC type in response to depolarizing steps from a holding potential of -70 mV. DB1 and FMB cells exhibited frequent, large spontaneous inhibitory postsynaptic currents (sIPSCs, Fig. 1A-B) that reversed near the calculated inhibitory reversal potential (see Fig 3B-C, G-H). DB3a cells showed less frequent sIPSCs, while tonic inhibitory events were infrequently observed in DB2 and DB3b cells. There was a significant difference in the current variance between cell types, reflecting in large part the difference in sIPSC frequency (Fig. 1C, one-way ANOVA, dF=4.0, F=8.65, p=3.2e-6). These results suggest that specific Off-CBC types receive more tonic inhibition than others. Similar tonic inhibitory events have been observed in Off-CBC cells of other mammalian retinas and have been shown to be of glycinergic origin (Ivanova et al., 2006; Graydon et al., 2018). Our next goal was to test whether the observed sIPSCs were glycinergic, and to determine the circuit origin of these events (see schematic Fig. 2A).

**Figure 5.**
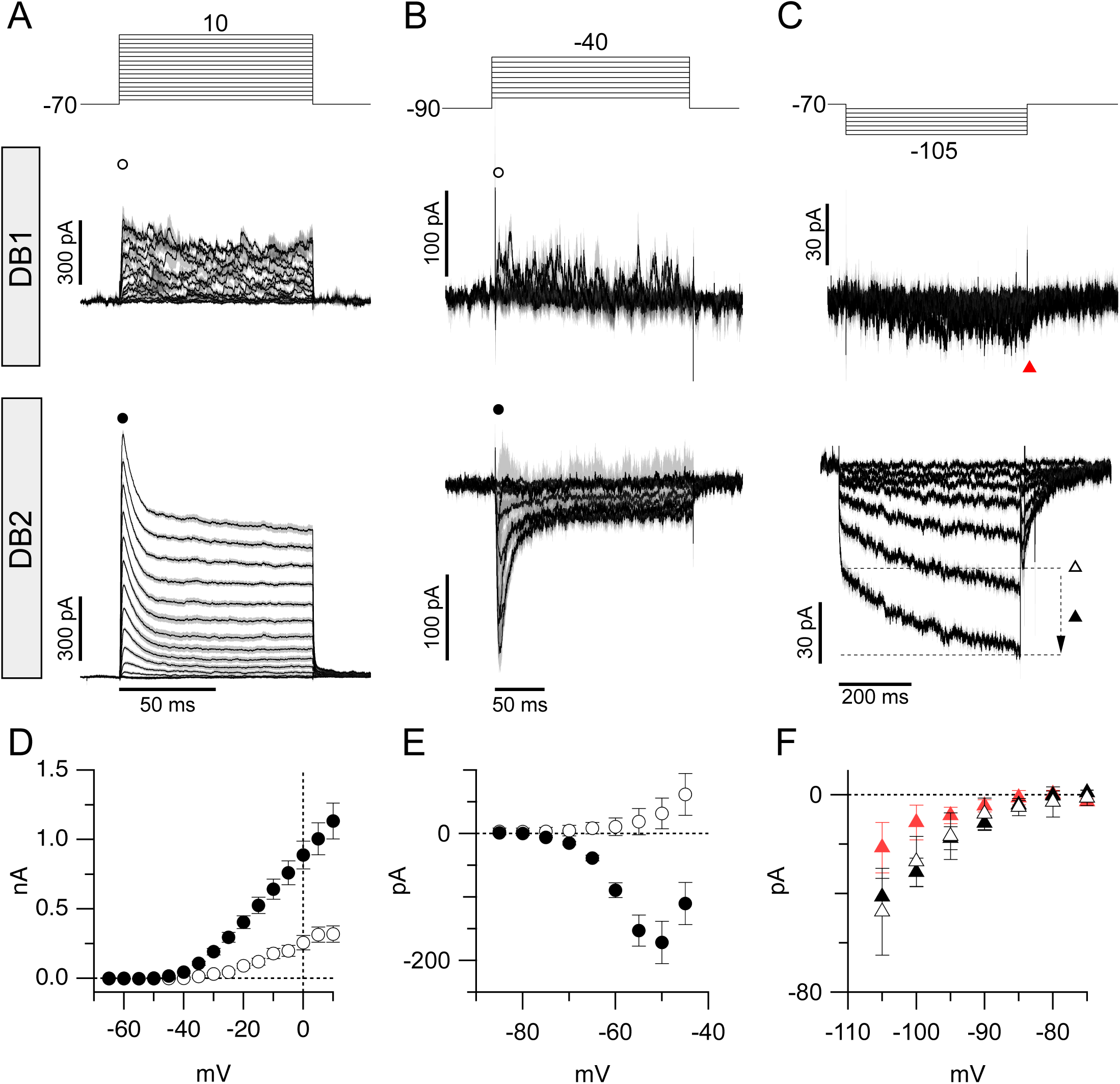
Voltage-gated currents in DB1 and DB2 bipolar cells. ***A-C***, Average leak-subtracted voltage-gated currents in DB1 and DB2 cells. Voltage step protocols are shown in top panels, and resulting currents from DB1 and DB2 cells are shown in the middle and bottom panels, respectively. Timing scale bar in lower panels applies to both cell types. Note prominent A-type potassium current, T-type calcium current and K_IR_ and I_h_ currents in DB2 cells that is absent in DB1 cells. Data are normalized to max amplitudes. Shading in A-C shows ± 1 s.e.m. ***D***, Comparison of peak outward potassium current in DB1 (open circles) and DB2 (solid circles) cells measured at a fixed timepoint indicated by the symbols (21.8 ms after onset of step). Measurement time points are indicated with corresponding symbols in A. ***E***, Comparison of inward current in DB1 (open circles) and DB2 (solid circles) cells at timepoints indicated by same symbols shown above traces in ***B*** (54 ms after step onset). ***F***, Comparison of instantaneous (open and red triangles) and time-dependent inward current (black triangles) in DB1 and DB2 cells. Instantaneous current amplitude was measured at a fixed time point (55.7 ms), time-dependent current is the difference between the minimum current amplitude and instantaneous current (black triangle). Data are mean ± s.e.m.

**Figure 6.**
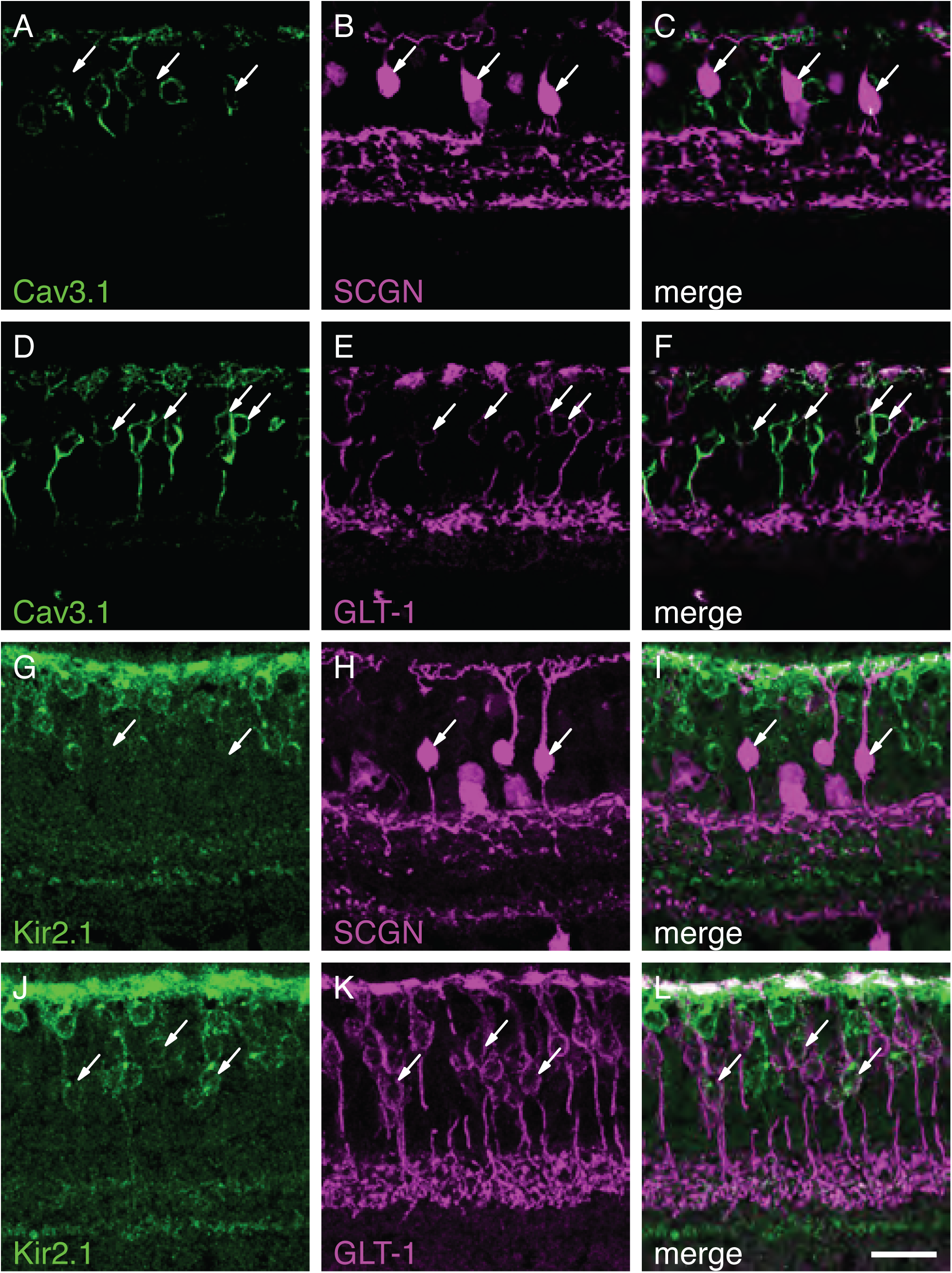
Voltage-gated ion channel subunits in primate DB1 and DB2 cells. ***A-F***, Localization of the T-type calcium channel subunit, Ca_V_3.1, in primate bipolar cells. Ca_V_3.1 is absent from SCGN+ DB1 cells (***A-C***, arrows). ***D-F***, A subset of GLT-1+ Off bipolar cells express Ca_V_3.1, consistent with expression in DB2 cells (arrows). ***G-L***, The inward-rectifying potassium channel (K_IR_) subunit, Kir2.1, is absent from DB1 cells (***G-I***, arrows) but present in the soma, dendrites and axon terminal boutons of GLT-1+ DB2 cells (***J-L***). Scale bar = 20 μm.

**Figure 7.**
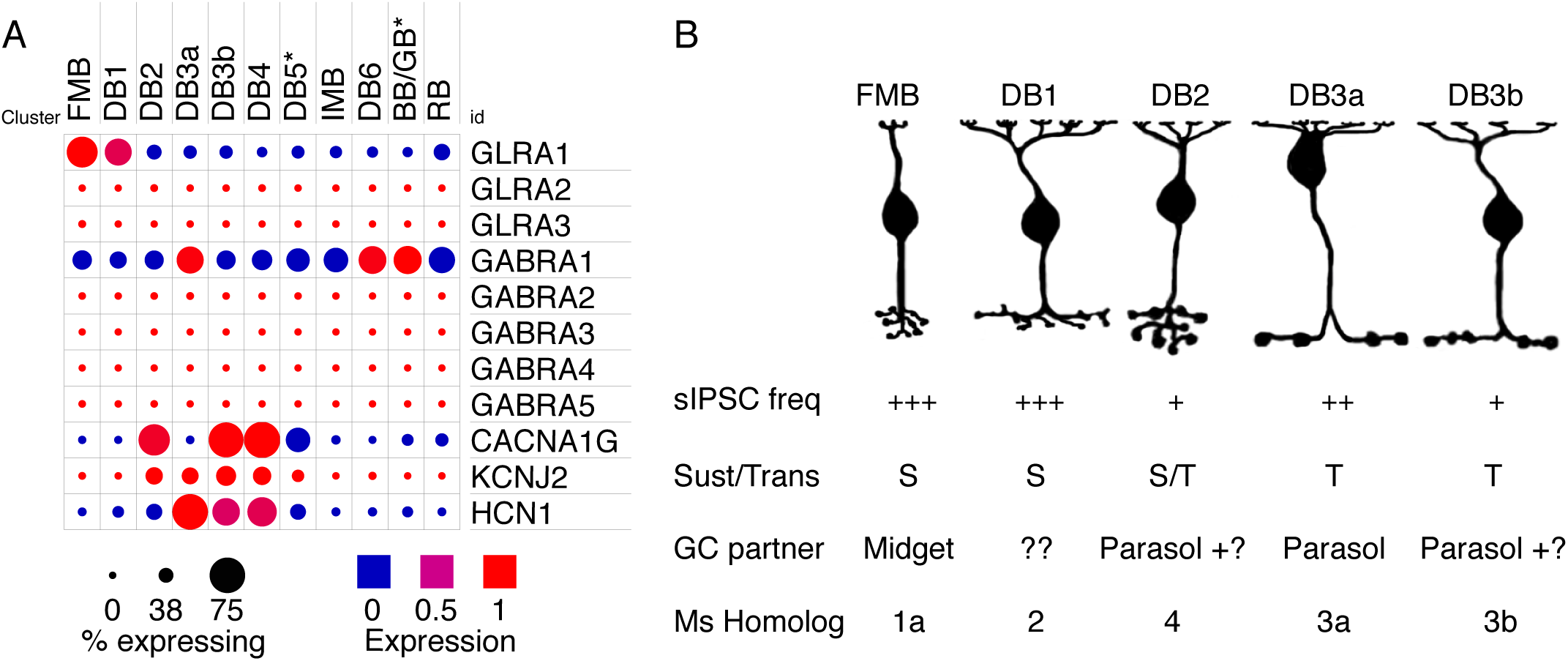
Summary of gene expression and physiological properties of macaque Off bipolar cells. ***A***, Dot-plots showing comparative gene expression of glycine receptor, GABA receptor and voltage-gated channel subunits in macaque peripheral bipolar cells. Dot color indicates expression level and dot diameter indicates the proportion of cells in the cluster in which the gene was detected. FMB and DB1 cells show highest levels of GLRA1 expression. Raw data from (Peng et al., 2019) plotted with the Broad Institute Single Cell Portal. ***B***, Summary of primate bipolar cell morphology, inhibitory inputs, physiological properties, ganglion cell (GC) partners and putative mouse bipolar homologs. Homology based on data from Peng et al., 2019.

### Spontaneous events are glycinergic and arise at the level of the axon terminals

Application of the glycine receptor antagonist, strychnine (0.5 μM) completely abolished the large sIPSCs in FMB cells (Figure 2B-C), consistent with a glycinergic input (Ctrl 19.1± 11.5 vs STR 0± 0 Hz, n=7 cells, mean ± s.d., p=0.004, Wilcoxon signed-rank test). The effect of strychnine was slowly reversible upon washout, and partial or complete recovery was achieved in 2 cells (Fig. 2C). If the sIPSCs arise from AII-ACs making glycinergic synaptic contacts on the axon terminals of Off-CBCs, then they should be absent in axotomized cells. To test this prediction, we recorded from midget bipolar cells in which axons had been cut during slice preparation (Fig. 2D). In the absence of an axon terminal, midget bipolar cells could be identified by their distinctive dendritic morphology because they contact a single cone pedicle. We also distinguished axonless On-midget bipolar (IMB) from Off-midget bipolar (FMB) cells by their responses to L-glutamate puffed onto their dendrites. FMB cells exhibited an inward current to L-glutamate application (Fig. 2D), consistent with activation of ionotropic glutamate receptors (Puthussery et al., 2014). None of the 5 axotomized FMB cells we recorded from exhibited sIPSCs, consistent with glycinergic inputs arising at the level of the axon terminals. Similar experiments were not performed in DB1 cells as these could not be unequivocally distinguished in their axotomized form and their sparsity precluded pharmacological experiments. Thus, we next tested for similarities in activation and kinetic properties of the observed sIPSCs in FMB and DB1 cells.

### Spontaneous IPSCs have kinetics consistent with alpha-1 subunit containing glycine receptors

To compare the properties of the sIPSCs in FMB and DB1 cells, we recorded sIPSCs at a range of holding potentials between -100 and 0 mV. Figure 3A shows sIPSCs in an FMB cell recorded at a range of holding potentials between -100 and 0 mV. At each potential, individual IPSCs were extracted using a matched template filter and averaged together to reveal the time-course and amplitude of the IPSCs as a function of membrane potential (Fig. 3B, see Materials & Methods). This analysis was applied to a group of 20 FMB cells to produce the average current-voltage relations shown in Fig. 3C/G. The mean sIPSC peak conductance for FMB cells was 0.97 nS and mean reversal potential was -76 mV, close to the calculated chloride reversal potential. The sIPSC rise-times were typical of synaptic events, with average 10-90% rise times ranging from 123-245 μs across the range of voltages tested (Fig. 3D). The decay phase of the sIPSCs was well fit by a single exponential function, with time constants exhibiting clear voltage dependence (Fig 3B,E, *τ* = 2.06± 0.24 ms at -100 mV vs 3.99± 0.21 ms at 0 mV, n = 20, mean ± s.e.m. t-test, p<0.0001). Similar analyses were conducted in 7 DB1 cells revealing sIPSCs largely comparable to that of the FMB cells (Fig. 3F-J). For DB1 cells, the average conductance was 0.80 nS and average reversal potential was -73 mV (Fig 3H). sIPSC rise times ranged from 206-278 μs and, like FMB cells, event decay time was voltage dependent (Fig. 3G,J, *τ* = 2.71± 0.2 ms at -100 mV vs 5.31± 0.25 ms at 0 mV (n=7 cells), t-test, p<0.0001). Taken together, the rapid kinetics and voltage dependence of the decay the spontaneous glycinergic events in FMB and DB1 cells are consistent with the presence of GlyRα1 containing receptors, in accord with findings in Off-CBCs from other species (Gill et al., 2006; Ivanova et al., 2006).

### Glycinergic input to Off-cone bipolar cells is driven by rod bipolar cells

If the glycinergic sIPSCs in Off-CBCs arise from the AII-ACs due to activation of the On-pathway, their frequency should be modulated by activation or blockade of the mGluR6 receptors expressed by rod bipolar cells and On-CBCs. To test this prediction, we bath applied L-AP4, an mGluR6 agonist that blocks tonic depolarization of rod bipolar and On-CBCs and, by extension, the AII-ACs (Slaughter and Miller, 1983). Application of L-AP4 reduced the frequency of the sIPSCs in FMB cells by 93% (Fig. 4A-B, Ctrl 19.2± 1.3 vs L-AP4 1.4± 1.5 Hz (n=4 cells), mean± s.d., p = 0.028, Wilcoxon test, partial washout in n=1 cell). Furthermore, puffing the mGluR6 receptor antagonist, CPPG, onto the OPL in the presence of L-AP4, to depolarize On-bipolar cells (Puthussery et al., 2009; Snellman et al., 2009), increased sIPSC frequency in FMB and DB1 cells and, to a lesser extent, DB3a cells (Fig. 4C-D). By contrast, similar CPPG puffs failed to elicit IPSCs in DB2 and DB3b cells (Fig. 4C-D), consistent with a lack of glycinergic input from AII-ACs. Together, these results indicate that glycinergic sIPSCs in Off CBCs arise through crossover inhibition driven by the On-pathway. These results further support the findings in Figure 1, suggesting that AII-AC-mediated glycinergic inhibition is directed primarily towards FMB and DB1 cells.

Since mGluR6 receptors are present on the dendrites of both rod bipolar and On-CBCs, the previous experiments do not determine which of these cell types drives glycinergic output from AII-ACs to Off-CBCs. Depolarizing On-bipolar cells with CPPG could depolarize the AII-ACs either through gap-junctions with On-CBCs, or via the glutamatergic synapse from rod-bipolar cells (Ghosh et al., 2001; Singer and Diamond, 2003). To determine whether rod bipolar cells represent a major component of the AII-AC mediated input to Off-bipolar cells, we blocked glutamatergic transmission at the rod bipolar to AII-AC synapse (Fig. 2A) using the AMPA receptor antagonist, GYKI 53655 (10 μM). On average, bath application of GYKI 53655 reduced the frequency of mIPSCs in FMB/DB1 cells by 81% (Ctrl 17.5± 6.1 vs GYKI 3.6± 1.4 Hz, n=5 FMB cells and n=1 DB1 cells, mean± s.d., p = 0.016, Wilcoxon rank test, partial washout in n=2 FMB cells, Fig. 4E-F). Thus, the sIPSCs observed in FMB cells are driven principally by the RBC → AII pathway, with a minor additional contribution from On-CBCs.

### Voltage-gated currents distinguish macaque Off-CBC types

The results presented thus far indicate that the glycinergic output of AII-ACs is directed towards specific Off-CBC types Our next objective was to determine the intrinsic functional properties of the Off-CBC types that receive AII-AC input. In a prior study, we showed that the transient, diffuse bipolar cell types DB3a, DB3b and the sustained flat midget bipolar (FMB) cells, have distinct inventories of voltage-gated currents that shape their response properties (Puthussery et al., 2013). Here, we measured the inventory of voltage-gated channels in DB1 and DB2 cells to determine whether their intrinsic properties favored transient or sustained signaling. First, we measured voltage-gated currents in DB1 and DB2 cells in response to depolarizing steps from -70 mV (Fig 5A, D). DB2 cells displayed a large (>1 nA) A-type outward potassium current (I_KA_), which distinguished them from DB1 cells, which had smaller and relatively sustained I_K_ (peak amplitude at +10 mV (mean ± s.d.); DB1; 318 ± 196 pA, n=11 cells vs DB2; 1132 ± 567 pA, n=19 cells, Welch’s t-test, p<0.0001). The large magnitude of I_KA_ also distinguishes DB2 cells from all other Off-CBC types described previously (Puthussery et al., 2013).

From a holding potential of -70mV, there was little evidence for voltage-gated sodium or calcium currents during depolarizing steps in either DB1 or DB2 cells (Fig 5A). However, since both of these channel types would be expected to show significant steady-state inactivation at -70 mV, we repeated the measurements from a holding potential of -90 mV, which is expected to relieve any such inactivation (Perez-Reyes, 2003; Puthussery et al., 2013) (Fig 5B, E). Under these conditions, DB2 cells exhibited a large, transient inward current that activated around -70 mV and reached an average peak amplitude of 171 ± 115 pA at ∼-50 mV (mean ± s.d., n=13 cells). A Boltzmann fit to the average I-V relation showed a V_1/2_ activation of -66 mV, rate (k) of 3.6 mV and maximal conductance of 2.64 nS. The activation kinetics (10-90% rise time 1.88 ± 0.54 ms, measured at -50 mV, mean ± s.d., n = 12 cells), time-course of inactivation (single exponential fit, decay *τ*, 6.67 ± 1.9 ms, mean ± s.d., n = 12 cells) and rate of recovery from steady-state inactivation (double exponential fit, fast phase, *τ* 1= 166 ± 33 ms, mean ± s.d., n = 3 cells) were consistent with properties of T-type Ca_V_ channels, specifically the Ca_V_3.1 subunit ((Perez-Reyes, 2003; Puthussery et al., 2013)). Similar inward currents were absent from DB1 cells, indicating that they do not express voltage-gated sodium or T-type Ca_V_ channels.

We previously showed that inwardly-rectifying potassium (K_IR_) and hyperpolarization-activated currents are present in transient (DB3a and DB3b), but not sustained (FMB), macaque bipolar cells (Puthussery et al., 2013). We tested for the presence of these currents in DB1 and DB2 cells by applying hyperpolarizing voltage steps from -70 mV (Fig. 5C, F). DB1 cells displayed small slowly-activating inward currents during hyperpolarizing steps, with a maximum amplitude of -21± 10 pA (n=9) (Fig. 5C,F). By contrast, DB2 cells displayed two non-linear inward current components of approximately equal magnitude; an instantaneous component (−47± 17 pA, n=13, solid symbols) and a voltage-gated component (−41± 27 pA, n = 13, open symbols), consistent with the presence of K_IR_ and I_h_, respectively. Overall, these results support the idea that DB1 cells have relatively passive intrinsic properties, whereas DB2 cells exhibit active conductances, expected to make light responses more transient (band-pass).

### Identity and localization of voltage-gated ion channel subunits in DB1 and DB2 cells

From our functional experiments, we found that DB2 cells exhibit Ih, Kir and T-type calcium currents, whereas these currents were absent in DB1 cells. To determine the molecular basis of these differences, we used immunohistochemistry to identify and localize the ion channel subunits underlying these currents (Fig. 6). In accord with our functional data, the Ca_V_3.1 subunit was localized to the somata of a subset of GLT-1 positive bipolar cells, but was absent from DB1 cells, which were identified with an antibody for secretagogin (SCGN) (Puthussery et al., 2011) (Fig. 6A-F). GLT-1 is present in both DB2 and FMB bipolar cells (Grünert et al., 1994), however, given the absence of T-type calcium currents in FMB (Puthussery et al., 2013) cells, it is likely that the DB2 cells express the Ca_V_3.1 subunit. We also sought to determine the origin of the inward currents in DB2 cells (Fig 6G-L). We found the inward-rectifier potassium channel subunit, Kir2.1, in DB2 cells, but not in DB1, cells. This channel subunit was abundant in the dendrites and present to a lesser extent in the soma, axon and axon terminal boutons. These molecular differences are further supported by transcriptomic data showing higher levels of expression of CACNA1G and KCNJ2 in DB2 compared with DB1 cells (Fig. 7A).

Together, the voltage-gated currents and immunohistochemical data shown here, and in our previous study (Puthussery et al., 2013), allow us to distinguish five primate Off-CBC types based on the amplitude and kinetics of I_K_, as well as the presence or absence of I_Na_, T-type I_Ca_, I_h_ and K_IR_ currents. Of these five types, DB1 and FMB were most similar with respect to their inventory of voltage-gated channels, but they can be easily distinguished by their characteristic dendritic and axonal morphologies. Our results support the idea that bipolar cells with axon terminals in stratum 1 of the IPL (FMB and DB1 cells) have more sustained physiological responses and relatively passive membrane conductances. Conversely, bipolar cells with axon terminals in stratum 2 of the IPL (DB2, DB3a and DB3b) have a variety of active conductances (T-type I_Ca_, I_Na_ and I_h_) that likely contribute to their transient physiological response properties.

Taken together, our results suggest that the glycinergic output of AII-ACs is biased towards sustained Off-CBC types. The functional properties of the glycinergic IPSCs are consistent with expression of the α1 glycine receptor subunit. Single-cell RNA sequencing data provide additional support for the conclusion that there is markedly higher expression of α1 glycine receptors (GLRA1) in DB1 and FMB cells than in other Off-CBC types (Fig. 7A).

## Discussion

A hallmark of the primary rod pathway is neural convergence that increases the sensitivity of ganglion cells at scotopic light levels by collecting signals from hundreds of rods. The glycinergic AII-AC represents the critical hub in the rod circuit, as they pass rod signals to On- and Off-type CBCs. Evidence from other mammals suggests that these connections are type-selective with some CBCs receiving minimal AII-AC input (McGuire et al., 1984; Tsukamoto and Omi, 2017; Graydon et al., 2018), consistent with the low scotopic sensitivity of some mouse and rabbit ganglion cells (Völgyi et al., 2004). Here, we have shown that the primate retina exhibits similar heterogeneity in primary rod pathway connectivity. Our results therefore predict that in primate retina, sustained type Off-CBCs should have higher scotopic sensitivity, and greater susceptibility to crossover inhibition under photopic light levels, than transient Off-CBC types.

### The primary rod pathway utilizes specific Off-CBC circuits

We found that specific Off-CBC types receive high levels of tonic inhibitory input that could be abolished by blocking glycine receptors, and strongly suppressed by blocking On-bipolar signaling or the rod bipolar cell to AII-AC synapse (Fig. 2A). These results indicate that glycinergic input to Off-CBCs arises from the well-described primary rod pathway (rod→ rod bipolar cell→ AII-AC→Off CBC). The sustained Off-CBC types, FMB and DB1, received the highest levels of tonic inhibition, whereas the transient types, DB2 and DB3b, received the least (summarized in Fig 7A). Our results are consistent with findings in mouse that show a bias in AII-AC output towards Type 2 Off-CBC (CBC2) (50-69%), and Type 3b (CBC3b) cells (17-19%) (Tsukamoto and Omi, 2013; Graydon et al., 2018). Functional studies in mouse further support the idea of selective AII-AC to Off-CBC connections (Mazade and Eggers, 2013). Mouse Type 1a and 2 bipolar cells, the likely homologs of FMB and DB1 cells (Fig. 7B, (Peng et al., 2019)) stratify in S1 of the IPL, and likely have relatively sustained physiology (Baden et al., 2013; Borghuis et al., 2013; Franke et al., 2017). A similar bias is apparent in cat retina, where S1-stratifying CBa1 Off-CBCs cells are the major recipient of AII-AC input (McGuire et al., 1984). Overall, our results in macaque, taken together with those from other mammals, strongly suggest a conserved motif, where AII-AC outputs are primarily targeted towards the sustained CBC pathways.

A further pattern that is emerging is that each Off-CBC type primarily receives primary or tertiary rod pathway input, but not both (Demb and Singer, 2012). For example, the mouse bipolar types, 3a, 3b and 4 make direct synaptic contact with rods (Mataruga et al., 2007; Haverkamp et al., 2008), but receive little AII-AC input. Our results align with these findings, in that DB3b cells, which showed the lowest levels of AII-AC input, are the only Off-CBC type to synapse directly with rods (Tsukamoto and Omi, 2014). Notably, in macaque, direct rod-to-Off-CBC input occurs less frequently than in mouse, a result that may explain the lower contribution of the tertiary rod pathway to scotopic signaling in primates (Grimes et al., 2018a).

### Glycinergic input to Off-CBCs is mediated by α1 glycine receptors

Glycine receptors (GlyRs) are comprised of one of four α subunits (α1, α2, α3, or α4) together with a modulatory β subunit (Lynch, 2009). The decay kinetics of α1 containing receptors are faster than those containing α2 or α3 (Singer et al., 1998). The rapid activation (∼200-300 μs) and decay kinetics (2-5 ms) of the glycinergic sIPSCs we recorded in Off-CBCs are consistent with the fast kinetics of α1β receptors and similar to glycinergic IPSCs in Off-CBCs of other mammals (Ivanova et al., 2006; Schubert et al., 2008; Wässle et al., 2009; Eggers and Lukasiewicz, 2011; Graydon et al., 2018). The GlyRα1 subunit is concentrated in the Off sublamina of the IPL, where it has been localized to contacts between AII-ACs and Off-CBCs (Sassoe-Pognetto et al., 1994; Grünert and Wässle, 2006). Indeed, in macaque retina, GlyRα1 puncta are localized at sites where DB1 and FMB cells contact AII-AC lobular appendages, and similar puncta have been reported to a lesser extent at DB3a axon terminals (Jusuf et al., 2005; Puthussery et al., 2011). The higher expression of GLRA1 (GlyRα1) in FMB and DB1 cells (Fig. 7A) compared to other Off-CBCs, provides additional evidence for biased output of AII-ACs towards FMB and DB1 cells, especially since other GlyR subunits are not detected at high levels in Off CBCs (Fig. 7A, raw data from Peng et al., 2019). Taken together, our data are consistent with expression of GlyRα1-containing glycine receptors at synapses between AII-ACs and FMB and DB1 cells in macaque retina.

### Voltage-gated channels in sustained versus transient Off-CBCs

We previously showed that CBCs that drive Off-midget and Off-parasol ganglion cells differ in their inventory of ion channels (Puthussery et al., 2013). Bipolar cells that provide input to parasol ganglion cells had prominent Na_V_ or T-type Ca_V_ currents, K_IR_ and prominent I_h_ currents, which contribute to their transient temporal response properties. Conversely, FMB cells, which provide excitatory input to midget ganglion cells, were relatively devoid of these channels. The results here extend on our previous analyses – DB2 cells, which elaborate their axon terminals in S2 of the IPL showed prominent T-type Ca_V_ currents and corresponding expression of Ca_V_3.1, whereas the S1-stratifying DB1 cells (Puthussery et al., 2011), lack these channels. Similarly, K_IR_ currents and the corresponding channel subunit, Kir2.1, were evident in DB2, but not DB1 cells. These data further support the idea that specific voltage-dependent ion channels contribute to the response properties of sustained vs transient bipolar cell types. T-type Ca_V_, I_h_ and K_IR_ channels are associated with CBCs that stratify near the middle of the IPL (Puthussery et al., 2013), findings further corroborated by data from single-cell sequencing (see Fig. 7A, (Peng et al., 2019)). Overall, our results indicate that signals transmitted through the primary rod pathway are directed towards Off-CBCs that are tuned to lower temporal frequencies. Primate rod signals are slowest under low scotopic conditions and speed up as background illumination increases (Grimes et al., 2018a). Thus, selectively routing rod signals into bipolar cells with low-pass temporal tuning would serve to preserve the fidelity of slow rod signals near scotopic threshold.

### Implications for scotopic signaling in the magnocellular and parvocellular pathway

Do all primate ganglion cells participate in scotopic vision? Or are some ganglion cell circuits exclusively driven by cone signals? Both Off-midget (sustained, parvocellular) and Off-parasol (transient, magnocellular) ganglion cells receive scotopic input (Lee et al., 1997; Field et al., 2009; Ala-Laurila and Rieke, 2014; Grimes et al., 2018a). Indeed, scotopic input to Off parasol cells was strongly suppressed by pharmacological blockade of the On-pathway (see Fig. 6E in Grimes et al, 2018a), indicating these inputs arise from the primary rod pathway. DB3a likely provide the major glutamatergic input to Off-parasol cells (Jacoby and Marshak, 2000). Although we showed DB3a cells receive some On-pathway driven inhibition (Fig. 4C-D), due to the sparsity of these cells we could not unequivocally determine the pharmacological basis of this inhibition. One possibility is that the On-pathway driven inhibitory events in these cells are mediated by GABA receptors (Figs, 3 and 6). Indeed, single-cell RNA sequencing data suggest that DB3a cells express high levels of GABRA1 (GABA_A1_), but lower levels of GLRA1 (GlyRα1) than DB1 and FMB cells (Fig. 7B). Although anatomical studies suggest that DB3a cells may receive some direct input from AII-ACs (Jusuf et al., 2005; Grünert and Wässle, 2006), rod signals may also be routed directly from AII-ACs to Off-parasol GCs. Indeed, such synaptic connections have been observed ultrastructurally (Bordt et al., 2006), and in mouse retina, rod signals are routed directly from AII-ACs to Off GCs at scotopic threshold (Arman and Sampath, 2012). Primate Off-parasol ganglion cells express GlyRα1 (Lin et al., 2000; Peng et al., 2019) and thus it is possible that they too receive scotopic input directly from AII-ACs. Pharmacological data suggests that Off parasol cells do receive glycinergic input driven by the On-pathway (Crook et al., 2014). Further studies are needed, however, to clarify the circuit that transmits rod input to Off-parasol ganglion cells.

We found that DB2 and DB3b cells receive little input from AII-ACs raising the possibility that the Off-type GCs downstream of these bipolar cells may receive less scotopic input than other ganglion cell types. In mice, scotopic sensitivity varies across Off-GC types and at least some Off-GCs are unaffected by blockade of the primary rod pathway (Völgyi et al., 2004). Here, we found that DB3b cells showed the lowest levels of glycinergic input. Though these cells make some synaptic input to Off-parasol cells, they presumably also drive other, as yet unidentified, ganglion cell types. Further electron microscopy reconstructions and circuit analyses will be required to quantify AII-ACs output to different primate Off-CBCs, and to establish the post-synaptic partners of the various Off-CBCs. Functional recordings from different types of primate Off-ganglion cell would also clarify whether specific primate output pathways are scotopically blind or exhibit lower scotopic sensitivity.

## Acknowledgments

We thank Rowland Taylor and Manoj Kulkarni for helpful comments and Rowland Taylor for sharing mini analysis scripts. This work was supported by NIH EY024265 (T.P.) and P30 Core Grant EY003176. Confocal imaging was conducted at the Molecular Imaging Center at UC Berkeley and supported by the Helen Wills Neuroscience Institute. We thank Holly Aaron and Feather Ives for microscopy training and assistance. Primate tissues were obtained with support of NIH P51OD011092 (ONPRC) and NIH P51OD011107 (CNPRC). K. Percival current address is Knight Cancer Institute, Oregon Health & Science University, Portland, OR, 97239.

